# Bento: A toolkit for subcellular analysis of spatial transcriptomics data

**DOI:** 10.1101/2022.06.10.495510

**Authors:** Clarence K. Mah, Noorsher Ahmed, Nicole Lopez, Dylan Lam, Alexander Monell, Colin Kern, Yuanyuan Han, Gino Prasad, Anthony J. Cesnik, Emma Lundberg, Quan Zhu, Hannah Carter, Gene W. Yeo

## Abstract

The spatial organization of molecules in a cell is essential for performing their functions. Spatial transcriptomics technologies have opened the door to characterization of cellular and subcellular organization. While current computational methods focus on discerning tissue architecture, cell-cell interactions and spatial expression patterns, these approaches are limited to investigating spatial variation at the multicellular scale. We present Bento, a Python toolkit that fully takes advantage of single-molecule information to enable spatial analysis at the subcellular scale. Bento ingests molecular coordinates and segmentation boundaries to perform three fundamental analyses: defining subcellular domains, annotating localization patterns, and quantifying gene-gene colocalization. To demonstrate the toolkit, we apply these methods to a variety of datasets including U2-OS cells (MERFISH), 3T3 cells (seqFISH+), and treated cardiomyocytes (Molecular Cartography). We quantify RNA localization changes in cardiomyocytes identifying mRNA depletion of critical cardiac disease-associated genes RBM20 and CACNB2 from the endoplasmic reticulum upon doxorubicin treatment. The Bento package is a member of the open-source Scverse ecosystem, enabling integration with other single-cell omics analysis tools.

## Introduction

The spatial organization of molecules in a cell is essential for performing their functions. While protein localization^1^ and disease-associated mislocalization are well appreciated^2, 3^, the same principles for RNA have begun to emerge. For instance, the spatial and temporal regulation of RNA play a crucial role in localized cellular processes such as cell migration and cell division^4, 5^, as well as specialized cell functionalities like synaptic plasticity^6–8^. Mislocalization of RNA has been associated with diseases such as Huntington’s disease (HD), where defects in axonal mRNA transport and subsequent translation in human spiny neurons lead to cell death and neurodegeneration^9–12^.

The study of subcellular RNA localization necessitates single-molecule measurements. Since the development of single-molecule fluorescent *in situ* hybridization (smFISH), recent advances in multiplexed methods such as MERFISH^13^, seqFISH+^14^, HybISS^15^, and Ex-Seq^16^ have enabled RNA localization measurements at near transcriptome scales, while maintaining single-molecule resolution. A number of computational toolkits, such as Squidpy^17^, stLearn^18^, Giotto^17^, Seurat^19^, and Scanpy^20^ enabled the characterization of tissue architecture, cell-cell interactions, and spatial expression patterns. Despite the single-molecule measurements in spatial transcriptomics, these analytical approaches are limited to investigating spatial variation at the multicellular scale and lack the ability to investigate subcellular organization. To further our understanding of RNA localization and its function in normal and abnormal cell activity, we need to expand our analytical capacity to the subcellular scale.

Recent methods such as FISH-quant-v2^21^ and FISHFactor^22^ identify subcellular patterns describing the spatial distribution of RNA species^22, 23^, but are constrained by the size of the data i.e. by the number of genes (1-100s), number of molecules (∼20,000) and do not scale to entire spatial transcriptomics datasets. Additionally, methods such as ClusterMap^24^ and Baysor^25^ highlight that subcellular spatial variation in transcript locations can infer cell and nucleus boundaries. In spatial proteomics data, CAMPA^26^ and Pixie^27^ utilize subcellular spatial variation in protein abundance to identify subcellular regions and annotate pixel-level features.

Building on these promising approaches, we present Bento, an open-source Python toolkit for scalable analysis of spatial transcriptomics data at the subcellular resolution. Bento ingests single-molecule resolution data and segmentation masks, utilizing geospatial tools (GeoPandas^28^, Rasterio^29^) for spatial analysis of molecular imaging data, and data science tools including SciPy^30^, and Tensorly^31^ for scalable analysis of high-dimensional feature matrices. Furthermore, Bento is a member of the Scverse ecosystem, enabling integration with Scanpy^20^, Squidpy^32^, and more than thirty other single-cell omics analysis tools.

## Results

### Overview of Bento data infrastructure for subcellular analysis

In order to facilitate a flexible workflow, Bento is generally compatible with molecule-level resolution spatial transcriptomics data (**Fig. 1A**), such as datasets produced by MERFISH^13^, seqFISH+^14^, CosMx (NanoString)^33^, Xenium (10x Genomics)^15, 34^, and Molecular Cartography (Resolve Biosciences)^35^. Bento’s

**Fig 1.**
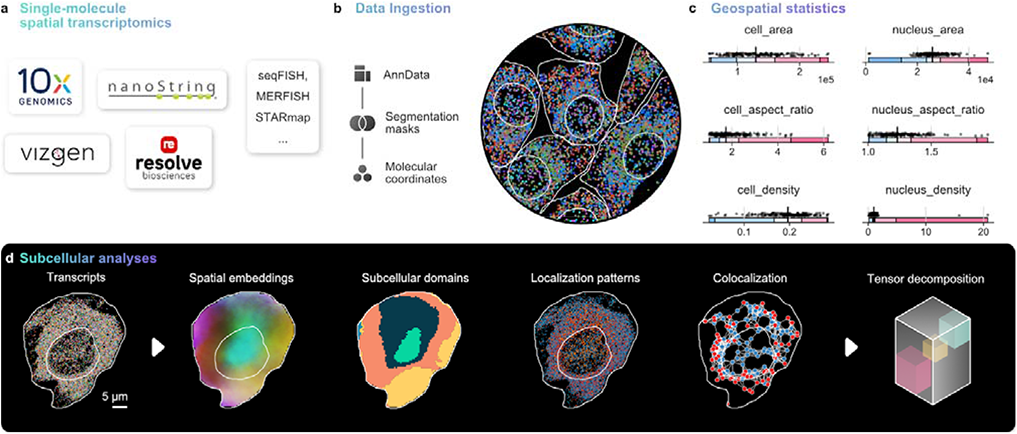
Workflow and functionality of Bento, subcellular spatial analysis toolkit. **A)** Single-molecule resolved spatial transcriptomics data from commercial or custom platforms can be ingested into Bento where it is converted to the AnnData format **(B)**, where it can be manipulated with Bento as well as a wide ecosystem of single-cell omics tools. **C)** Geometric statistics are illustrated for the seqFISH+ dataset, including metrics describing cell and nuclear geometries and cell density to assess overall data quality. **D)** Bento has a standard interface to perform a wide variety of subcellular analyses.

workflow takes as input 1) 2D spatial coordinates of transcripts annotated by gene, and 2) segmentation boundaries (e.g. cell membrane, nuclear membrane, and any other regions of interest) (**Fig. 1B**). If available, Bento can also handle arbitrary sets of segmentations for other subcellular structures or regions of interest. These inputs are stored in the AnnData data format^36^, which links cell and gene metadata to standard count matrices, providing compatibility with standard single-cell RNA-seq quality control and analysis tools in the Scverse ecosystem^20^. With a data structure for segmentation boundaries and transcript coordinates in place, Bento can easily compute spatial statistics and measure spatial phenotypes to build flexible multidimensional feature sets for exploratory subcellular analysis and utilize these spatial metrics to augment quality control (**Fig. 1C**).

Bento offers a precise yet flexible palette of subcellular analyses (**Fig. 1D**). First, we propose RNAflux, an unsupervised method for semantic segmentation of subcellular domains. For every pixel in a cell, pixels are first embedded by quantifying local gene composition. Clustering pixels in the embedding space identifies consistent subcellular domains that can be functionally characterized with known gene signatures. In contrast to unsupervised analysis, we additionally present RNAforest, a multilabel model for annotating RNA localization patterns adapted from FISH-quant v2^21^. RNAforest classifies RNA transcripts based on their localization with respect to landmarks (i.e. cell and nuclear boundaries). Finally, we implement an approach to systematically measure pairwise colocalization of transcripts in a compartment-specific manner. We demonstrate the utility of these methods by characterizing localization changes in human iPSC-derived cardiomyocytes upon treatment with doxorubicin, a widely used chemotherapeutic known to cause cardiotoxicity^37^.

### RNAflux: Unsupervised semantic segmentation of subcellular domains in single cells

RNAflux, quantifies spatial composition gradients within the bounds of a given area. The embedding is defined as the local RNA composition for a fixed radius **r**, normalized by the parent cell composition to emphasize intracellular variation and minimize cell to cell variation. By applying the embedding at regular intervals (e.g. every pixel), this generates a continuous spatial embedding across the area of every cell (**Fig. 2A, Methods**).

**Fig 2.**
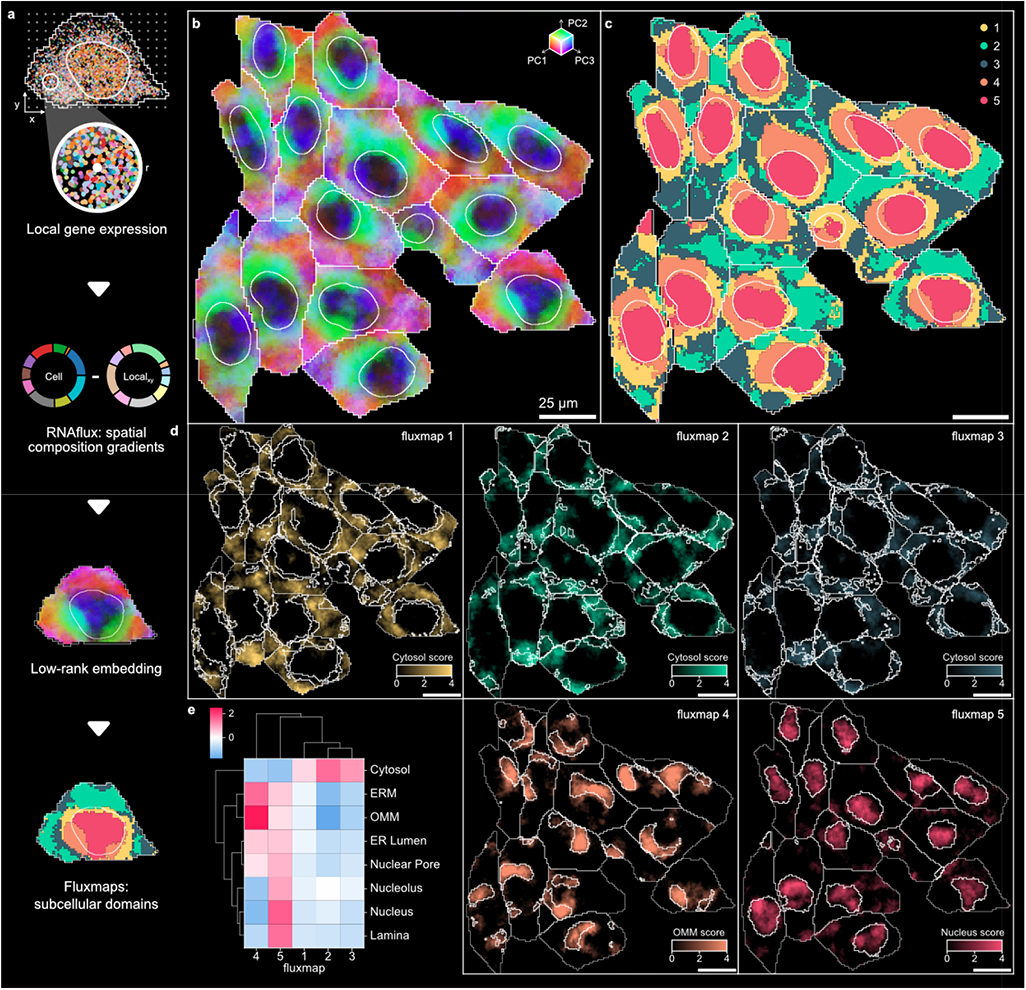
Unsupervised semantic segmentation of subcellular regions in single cells using RNAflux. **A)** Local neighborhoods of a defined radius are arrayed across a cell and a normalized gene composition is computed for each pixel coordinate, producing an RNAflux embedding. **B)** The first three principal components of the RNAflux embedding are visualized for U2-OS cells coloring RGB values by PC1, PC2, and PC3 values respectively for each pixel. **C)** Fluxmap domains are computed from each RNAflux embedding to create semantic segmentation masks of each subcellular domain. **D)** Relative enrichment of organelle-specific expression derived from APEX-seq data (Fazal et al. 2019) across RNAflux embedding space. The most highly enriched organelle is shown for each fluxmap domain. Domain boundaries are denoted by white lines within each cell. **E)** Mean enrichment score of pixels contained within each fluxmap domain. Pixel enrichment scores are defined as the weighted sum of a pixel’s RNAflux embedding and loadings for a given organelle-specific geneset (**Methods**).

Applied to a MERFISH dataset with a target panel of 130 genes in U2-OS cells, we demonstrate that RNAflux embeddings can detect meaningful spatial composition gradients strongly indicative of organelles despite the small number of genes. Performing dimensional reduction of the embeddings showed that the top sources of variation spatially correspond to the nucleus, the nuclear periphery, and cytoplasmic regions consistently across cells (**Fig. 2B, Methods**) confirming that RNAflux captures intracellular variation. To identify compositionally similar areas in a data-driven manner, we used self-organizing maps (SOMs) to perform unsupervised semantic segmentation of the embeddings (**Methods**). We denote the resulting clusters as “fluxmap domains”. This assigned pixels to 5 fluxmap domains highlighting spatially consistent regions across every cell (e.g. fluxmap 5 is found in the nucleus while the remaining domains constitute the cytoplasm) (**Fig. 2C**). Adjacent pixels tended to be assigned the same label, resulting in visually contiguous domains. We also embedded a seqFISH+ dataset of 3T3 cells with a panel of 10,000 genes, and were able to find similar fluxmap domains despite higher plexity and lower transcript detection efficiency (**Supp. Fig. 1A-1B**).

Finally, we sought to characterize the fluxmap domains with known information about organelle-specific gene expression. We used data from a previous study that captured organelle-specific expression via APEX-seq, a technique performing proximity labeling and sequencing of RNA^38^. The enrichment score for each pixel is calculated by taking the weighted sum of its RNAflux embedding and loadings for a given organelle-specific geneset (**Methods**). Visualizing each pixel’s organelle-specific enrichment scores from the APEX-seq dataset highlights the subcellular localization of these compartments, such as the cytosol, nucleus, nucleolus, nuclear pore, endoplasmic reticulum lumen (ER lumen), ER membrane (ERM), and the outer mitochondrial membrane (OMM) (**Fig. 2D**). We find the nuclear compartments have high scores in domain 5, while the cytoplasm scores rank highest in domains 1, 2, and 3. Both the ERM and OMM scores are the strongest in domain 4 (**Fig. 2E**). In contrast, organelle enrichment in the seqFISH+ dataset does not spatially correspond as well (**Supp. Fig. 1C**); non-intuitively, nucleus-specific enrichment highlights the cell periphery (**Supp. Fig. 1D**). It is known that capture efficiency is drastically lower in the nuclear region of this particular dataset, artificially lowering the local expression and potentially confounding the enrichment measure.

In summary, RNAflux finds subcellular regions that strongly correspond to organelles in terms of spatial organization and local gene composition. As an unsupervised method, RNAflux can be applied to any cell type for inferring subcellular domains from transcript locations.

### RNAforest: Utilizing subcellular landmarks to predict RNA subcellular localization

In computer vision, key points or landmarks are commonly used for tasks like facial recognition^39^ and object detection. Analogous to these classical applications, we derive spatial features using cell and nucleus boundaries as landmarks to predict RNA localization patterns from spatial summary statistics. Building on the summary statistics used for classifying smFISH data in FISH-quant v2^23^, RNAforest consists of an ensemble of five binary random forest classifiers rather than a single multi-classifier model to assign one or more labels. These pattern labels, adapted from several high-throughput smFISH imaging experiments in HeLa cells^40–43^, are broadly applicable to eukaryotic cells: (i) nuclear (contained in the volume of the nucleus), (ii) cytoplasmic (diffuse throughout the cytoplasm), (iii) nuclear edge (near the inner/outer nuclear membrane), (iv) cell edge (near the cell membrane), and (v) none (complete spatial randomness).

We used the FISH-quant simulation framework to generate realistic ground-truth data^42^. Each sample is defined as a set of points with coordinates in two dimensions, representing the set of observed transcripts for a gene in a particular cell. In total, we simulated 2,000 samples per class for a total of 10,000 samples (**Methods**). We used 80% of the simulated data for training and held out the remaining 20% for testing. Each sample is encoded by a set of 13 input features, describing characteristics of its spatial point distribution, including proximity to cellular compartments and extensions (features 1-3), measures of symmetry about a center of mass (features 4-6), and measures of dispersion and point density (feature 7-13) (**Fig. 3A**). These features are normalized to morphological properties of the cell to control for variability in cell shape. A detailed description of every feature is described in **Supp. Table 1**.

**Fig 3.**
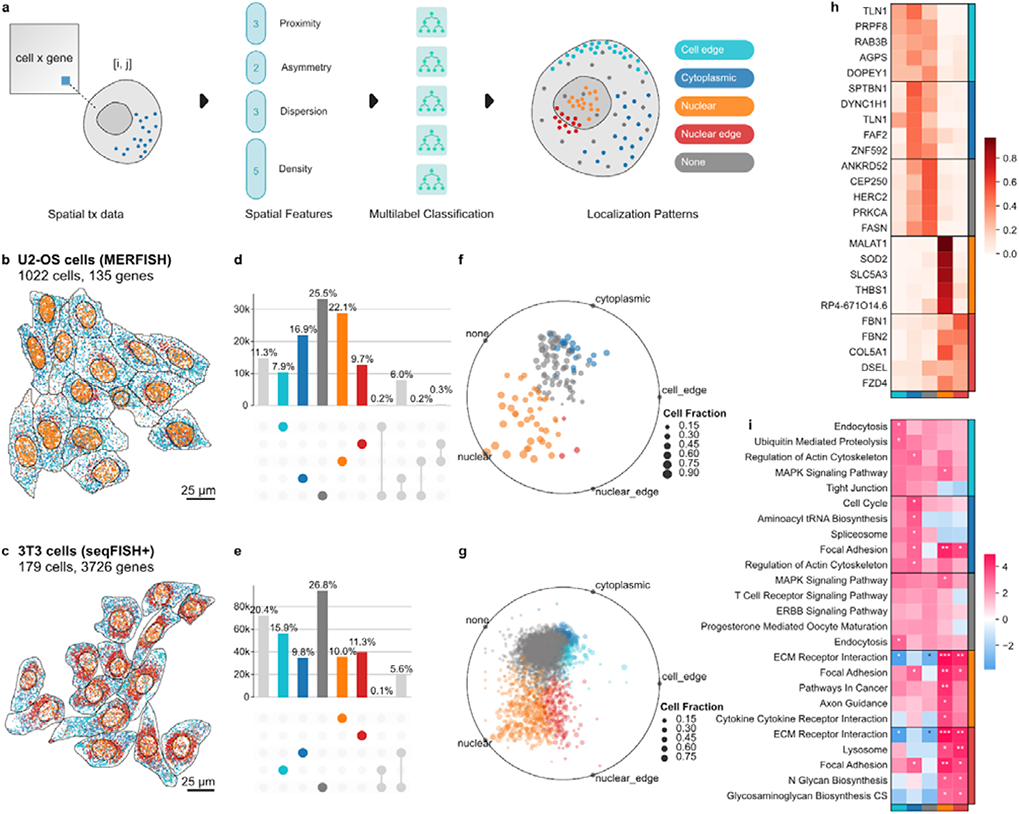
Subcellular localization patterns identified with RNAforest. **A)** Thirteen spatial summary statistics are computed for every gene-cell pair describing the spatial arrangement of molecules and boundaries in relation to one another. The features are inputs for RNAforest, a multilabel classifier which assigns one or more subcellular localization labels: cell edge, cytoplasmic, nuclear, nuclear edge, and none. Top 10 genes for each label visualized for each label other than “none” in **B)** U2-OS cells, and **C)** 3T3 cells. **D)** and **E)** show the proportion of measured transcripts assigned to each label. **F)** and **G)** show for each gene the relative label proportion across cells and is colored by the majority label (**F** and **G**). **H)** Top 5 consistent genes for each label. I) ssGEA identifies enrichment of GO cellular component domains for each label in the 3T3 cell dataset.

We applied RNAforest on the MERFISH dataset measuring 130 genes (low plexity) in U2-OS cells and high detection efficiency per gene (111 molecules per gene per cell on average), and on the seqFISH+ dataset measuring 10,000 genes (very high plexity) but lower detection efficiency (8 molecules per gene per cell on average) (**Fig. 3B-C, Supp. Fig. 2)**. In agreement with previous work characterizing RNA localization of 411 genes^43^, we find that genes commonly exhibit variability in localization across cells. This suggests that heterogeneity in localization likely generalizes to the entire transcriptome. Of the localization patterns besides “none”, “nuclear” was the most common (22.1%) in the U2-OS osteosarcoma cells (**Fig. 3D & 3F**), while “cell edge” was the most common (15.9%) in the 3T3 fibroblast cells (**Fig. 3E & 3G**).

In the U2-OS cells, we found many genes to have preferential localization in different subcellular compartments (**Fig. 3H**). We identify known nuclear localized genes such as MALAT1^44, 45^, as well as “nuclear edge” genes such as the ECM component genes FBN1, FBN2, and COL5A^13^. As expected, we find genes preferentially “nuclear” and “nuclear edge” localized to mirror nucleus and endoplasmic reticulum genes found in a 10k genes MERFISH study in U2-OS cells with the addition of ER staining^46^ (**Supp. Fig. 3, Methods**). Leveraging the 3T3 seqFISH+ dataset’s higher plexity, we were able to ask whether genes with similar localization preference are functionally related. We applied gene set enrichment analysis to gene localization frequencies to identify enriched gene ontology terms^47^ (**Fig. 3I, Methods**). Secr**e**tory processes were enriched in the nucleus and nuclear edge, which may be linked to increased transcription of fibroblast-related functions. Cell edge enriched pathways consisted of those with the cell membrane as their site of function (e.g. endocytosis and tight junction suggesting local translation of these genes). Additionally, the term for cell cycle was significantly enriched in the cytoplasm only. Genes without strong localization preference (most frequently “none”) were not significantly associated with any pathways. These genes likely do not undergo active transport and are functionally independent of local translation.

### Bento finds intermolecular interactions from compartment-specific colocalization

Prior spatial transcriptomics studies have used a number of colocalization metrics from the Geographic Information Systems and Ecology fields e.g. the univariate and bivariate versions of the Ripley’s K function (also known as cross-k-function)^48^, Moran’s I^49^, and the join count statistic^50^. For measured spatial associations to contribute to a biological understanding, the symmetry of the association is important to capture (e.g. if RNA A spatially correlates with RNA B, but the inverse is not necessarily true), especially when there are many more points in one set than the other. As such, in Bento we implemented the Colocation Quotient (CLQ)^51^, which given two sets of points, A and B, finds the ratio of observed to expected proportions of B among A’s neighbors (**Methods**). The CLQ can be understood as a modification of the cross-k-function that accounts for autocorrelation and asymmetry.

The spatial distribution of mRNA changes along various stages of the RNA life cycle, from pre-mRNAs to mature mRNAs. For example, transcriptional bursting can lead to strong colocalization of transcripts near the site of transcription, while transport of transcripts towards their eventual subcellular destinations may be more varied. Therefore, looking at pairwise colocalizations over the entire cell can limit the detection of spatial co-regulation, motivating the need for compartment-specific measurements. We quantified the differences in RNA colocalization in the nucleus versus the cytoplasm by calculating CLQ scores considering each compartment separately. Applied to every gene-gene pair across every cell produces a tensor of shape **P** x **C** x **S** where **P**, **C**, and **S** represent the number of gene-gene pairs, cells, and subcellular compartments, respectively (**Fig. 4A, Methods**).

**Fig 4.**
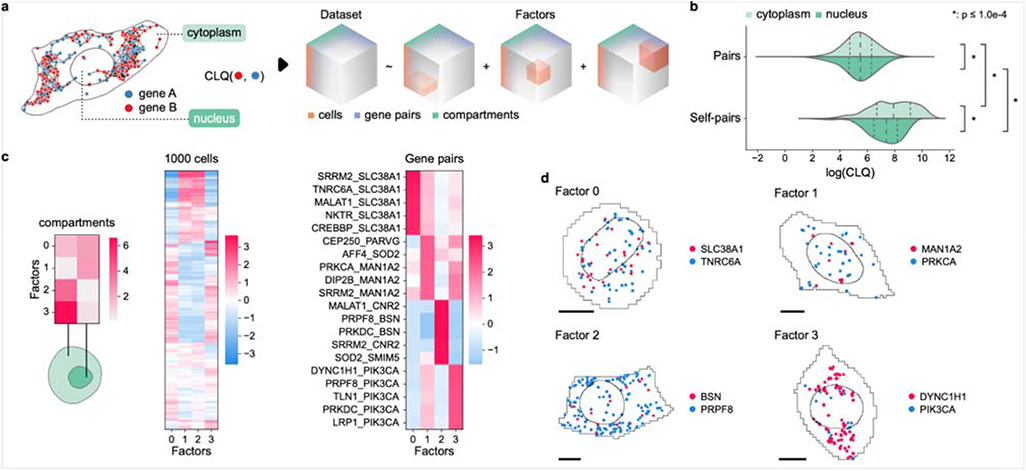
Compartment-specific colocalization of RNA. **A)** Transcripts are separated by compartment (nucleus and cytoplasm) before CLQ scores are calculated for every gene pair across all cells. This yields a cell x gene pair x compartment tensor. **B)** Comparison of log CLQ distributions for gene pairs and self-pairs, further categorized by each factor are shown. **D)** Top examples of compartment-specific colocalized gene pairs. Black scale bars denote 10 µm.

To quantify colocalization in a compartment-specific manner, we employed tensor decomposition — specifically, non-negative parallel factor analysis — a data-driven, unsupervised approach for discovering substructure in high-dimensional data^31, 52^. We applied tensor decomposition to decompose the U2-OS dataset colocalization tensor into **k = 4** “colocalization factors”. The number of factors was determined using the elbow method heuristic, optimizing for the root mean squared error (RMSE) reconstruction loss (**Methods**). Missing values are ignored when calculating the loss. Unlike matrix dimensionality reduction methods, such as PCA, the order of the components (factors) is unassociated with the amount of variance explained. Each of the 4 colocalization factors is composed of 3 loading vectors, which correspond to the compartments, cells and gene pairs. Higher values denote a stronger association with that factor. Crucially for interpretation, factors derived from tensor decomposition are not mutually exclusive and share overlapping sets of associated compartments, cells, and gene pairs.

An initial comparison of global colocalization between nuclear and cytoplasmic fractions unsurprisingly found that RNA transcripts tend to colocalize more strongly with themselves than with other RNA transcripts (**Fig. 4B**). Additionally, self-colocalization is significantly stronger in the cytoplasm than in the nucleus, suggesting that RNA transport is a transcriptome-wide mechanism of regulation and not limited to specific transcripts. Considering the colocalization factors, these trends are broken down into unique combinations of colocalization behavior (**Fig. 4C**). Factor 0 captures gene pairs in a subpopulation of cells that tend to colocalize across the entire cell, with pairs including SLC38A1 showing the strongest signal. Factor 3 describes gene pairs in mostly the same cell subpopulation, that colocalize specifically in the cytoplasm. Pairs including PIK3CA dominate this behavior. In the complementary cell population, Factors 1 and 2 highlight colocalized gene pairs in the nucleus and cytoplasm respectively. Strikingly, the top gene pair in factor 2 is MALAT1 and CNR2; MALAT1 is known to primarily localize to the nucleus, but here shows strong colocalization in the cytoplasm.

### Doxorubicin-induced stress in cardiomyocytes depletes RNA from the endoplasmic reticulum

Doxorubicin (DOX) was once one of the most effective broad-spectrum anti-cancer anthracycline antibiotics^53, 54^ with particular efficacy against solid malignancies such as lung and breast cancer, as well as hematologic neoplasia^55, 56^. However, DOX’s propensity to cause cardiac damage in patients has led to significant limitations in its clinical use^57^. The exact mechanism by which DOX induces heart failure is unclear, but significant evidence suggests cardiomyocyte injury driven by oxidative stress as a major factor^55, 58–61^. Specifically, DOX causes stress and dysfunction in multiple cellular compartments in cardiomyocytes such as mitochondria, Sarco/endoplasmic reticulum (SER), deficiencies in calcium signaling, and lipid degradation at the cellular membrane^62^. We reasoned that by measuring the localization of the RNA transcripts of 100 genes crucial to cardiomyocyte health and function (**Supp. Table 2**), and leveraging the tools developed within Bento, we could recapitulate known dysfunction of subcellular domains in cardiomyocytes upon DOX stress and measure novel RNA localization phenotypes.

We utilized a chemically-defined protocol to differentiate human induced pluripotent stem cells (iPSCs) into beating cardiomyocytes and treated them with either DMSO (vehicle) or 2.5 μM DOX for twelve hours before fixation (**Methods**). Single molecule spatial transcriptomes were measured by Resolve Bioscience using Molecular Cartography. The resulting data was segmented using ClusterMap^24^ for cell boundaries and Cellpose^63^ for nuclei boundaries (**Fig. 5A**). Non-myocytes were filtered out using SLC8A1 as a canonical marker for cardiomyocytes (**Methods, Supp. Fig. 4A**). Comparing vehicle and DOX treated cardiomyocytes, we found NPPA, a classic marker for cardiac stress^64, 65^, to be upregulated in DOX treated cells (**Fig. 5B**). We identified subcellular domains in vehicle and DOX treated cardiomyocytes using RNAflux, clustering the domains into four fluxmap domains (**Fig. 5C**). Enrichment of organelle-specific gene expression aligned domains to the nucleus (nuclear pore, nucleolus, and nucleus), ERM and OMM, ER lumen, and cytosol respectively (Fig. 5C & D, **Supp. Fig. 4C**). Comparing the gene composition in each domain, we observe an overall localization bias towards both the nucleus and ERM/OMM in vehicle treated cells (**Fig. 4E top**), in agreement to prior poly(A) smFISH studies^66^. However, RNA in the DOX treated cardiomyocytes demonstrated a shift in average RNA localization away from the ERM/OMM and towards the nucleus (**Fig. 5C bottom**). There is evidence that 90% of genes have a half life of less than 260 minutes^67^, far less than the 12 hour DOX treatment, indicating that the shift in RNA localization is likely due to reduced nuclear export of newly synthesized RNA from the nucleus to the ERM/OMM. Indeed, even low concentrations of DOX have been demonstrated to alter structural fibrous proteins as well as mitochondrial depolarization and fragmentation^68^. Of particular note, the RNA binding protein RBM20 – a critical regulator of mRNA splicing of genes encoding key structural proteins associated with cardiac development and function – had a pronounced depletion of RNA transcripts outside of the nucleus upon DOX treatment (**Fig. 5F**). With further validation, this may indicate nuclear retention and or degradation of nuclear exported RBM20 mRNA as a potential mechanism of DOX induced cardiomyopathy. Similarly, we found the mRNA of calcium voltage-gated channel subunit CACNB2 to also deplete outside of the nucleus (**Fig. 5G**). The loss of CACNB2 translation outside of the nucleus may impact calcium signaling crucial to cardiomyocyte function^69^.

**Fig 5.**
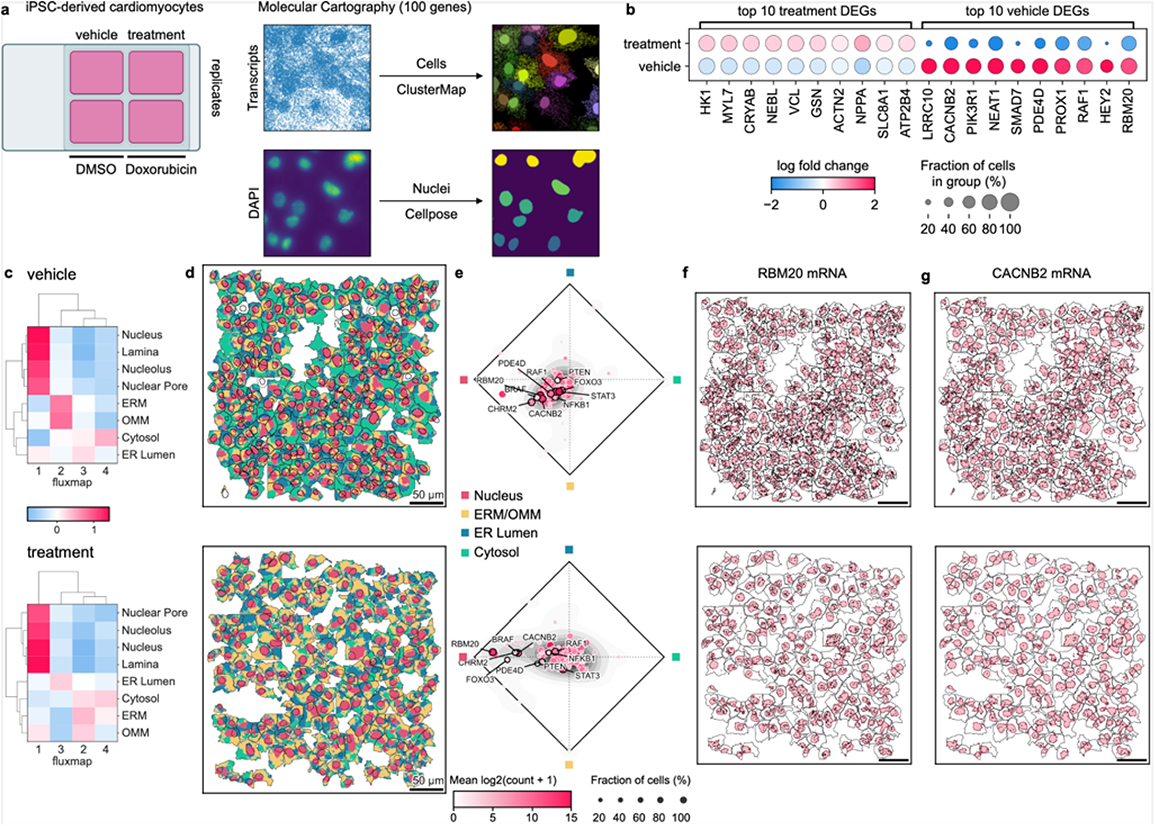
Subcellular RNA localization changes upon Doxorubicin treatment in iPSC-derived cardiomyocytes. **A)** Cardiomyocytes derived from human iPSCs were treated with DMSO or 2.5 μM DOX for 12 hours. The localizations of 100 genes relevant to cardiomyocyte health and function were measured using Molecular Cartography. Cell boundaries were determined using ClusterMap and nuclei were segmented using Cellpose. **B)** Top 10 upregulated and downregulated genes in vehicle versus treatment. **C)** Organelle-specific gene enrichment of fluxmap domains for the cytosol, endoplasmic reticulum membrane (ERM), endoplasmic reticulum lumen (ER Lumen), nuclear lamina, nucleus, nucleolus, nuclear pore and outer mitochondrial matrix (OMM). **D)** Fluxmap domains visualized for a representative field of view of cardiomyocytes for vehicle and treatment respectively highlighting cellular nuclei, ERM/OMM, ER Lumen, and cytosol. **E)** RNAflux fluxmap enrichment of each gene averaged across vehicle and treatment cardiomyocytes captures changes in subcellular RNA localization. Visualization of RBM20 (**F**) and CACNB2(**G**) confirms the depletion of transcripts from the perinuclear and cytosolic compartments of cardiomyocytes upon DOX treatment.

## Discussion

Bento seeks to interrogate subcellular biology via its “subcellular first” approach to spatial analysis, complementary to “cell-type or tissue first” spatial analysis methods. We implement novel methods to tackle 3 fundamental analyses: RNAflux for semantic segmentation of subcellular domains, RNAforest to identify mRNA localization patterns, and compartment-aware approach for colocalization analysis. We demonstrate Bento’s versatility by successfully applying it to two datasets from different technology platforms, detection efficiency, and image processing. In U2-OS cells, we use RNAflux to uncover seven distinct subcellular domains from a panel of 130 genes. Despite the fact that the gene panel was not designed to include organelle marker transcripts, RNAflux performed effectively, highlighting its potential for discovery and generalizability to all spatial transcriptomics experiments. By using RNAforest to annotate RNA localization patterns in both U2-OS and 3T3 cells, we find subcellular mRNA localization reflects gene function. We also explore the use of CLQ scores combined with tensor decomposition to identify pairwise colocalization patterns that are unique to the nuclear or cytoplasmic domains of U2-OS cells. To take it a step further, we applied Bento to quantify RNA localization changes in iPSC-derived cardiomyocytes with doxorubicin treatment. RNAflux identified four subcellular regions corresponding to the nucleus, ERM/OMM, ER Lumen, and Cytosol. We found DOX treated cardiomyocytes to be depleted of RNA in the endoplasmic reticulum, causing mRNA encoding important cardiac genes such as RBM20 and CACNB2 to remain in the nucleus.

Through the process of developing and iterating Bento and its tools such as RNAflux and RNAforest, we were able to reflect on RNA localization as well as the spatial transcriptomic methods used to measure it. A frequent trade-off comparing various spatial transcriptomics technologies is plexity versus detection efficiency. From the context of subcellular biology, each carries particular benefits that can be considered during experimental design. Most current commercial platforms for spatial transcriptomics use a substantial part of the gene panel – if not the entirety of it – for cell type markers. While this may be well suited for spatial single-cell atlasing, it substantially limits what is “interesting” from a subcellular standpoint, namely RNA that serve particular cellular functions within a cell. For this reason we fully customized our cardiomyocyte gene panel for Molecular Cartography to emphasize RNA localization and expression of cardiac functional genes.

Classically, the study of RNA has centered around its role as an intermediary information molecule encoding genomic information for protein synthesis. We began our investigation of RNA localization with the hope of understanding how the spatial organization of RNA functions as a mechanism for post-transcriptional regulation. As spatial transcriptomics grows in popularity, we hope Bento is a platform for the tools needed to quantify the complex molecular dynamics governing normal and abnormal cellular processes.

## Methods

### MERFISH and seqFISH+ data preprocessing

For the seqFISH+ dataset, we limited the scope of our analysis to the set of genes for which at least 10 molecules were detected in at least one cell. This helped reduce sparsity in the data, resulting in 3726 genes remaining. Because pattern classification requires nuclear segmentation masks, we removed all cells lacking annotated nuclei for a remainder of 179 cells. Because the MERFISH data had a much higher number of molecules detected per gene, no gene needed to be removed. Again, cells without annotated nuclei were removed, leaving 1022 cells for downstream analysis.

### Preprocessing cardiomyocytes datasets

Single-cell expression matrices of both vehicle replicates and both DOX treatment samples were concatenated as a single expression matrix. Cells were projected into two dimensions with UMAP dimensional reduction. No significant batch effects were detected. Leiden clustering was performed at resolution=0.5 to isolate and filter out a non-myocyte population depleted in SLC8A1 expression (**Supp. Fig. 4A**). All described preprocessing steps were performed in Scanpy^20^.

### RNAflux: Unsupervised spatial embedding and subcellular domain quantization

To generate RNAflux embeddings, first a set of query coordinates are generated tiling across the cell area on a uniform grid. This effectively downsamples the original data units (pixels) resulting in much fewer samples to compute embeddings. For the MERFISH U2-OS dataset, a step size of 10 data units (pixels) was used to generate the uniform grid. For the iPSC-derived cardiomyocytes, a step size of 5 data units was used. Each query point is assigned an expression vector, counting the abundance of each gene within a fixed radius of 50 data units. Each expression vector is normalized to sum to one, converting the expression vector to a composition vector. Similarly, the cell composition vector is calculated by normalizing the total cell expression to sum to one. The RNAflux embedding at a given query coordinate is defined as the difference between the query composition and its corresponding cell composition, divided by the standard deviation of each feature within each cell.

The RNAflux embedding serves as an interpretable spatial gene embedding that quantifies highly local fluctuations in gene composition. Dimensional reduction of the embeddings is performed using truncated singular value decomposition (SVD). Truncated SVD was chosen over PCA to better handle large but sparse data. Embeddings were reduced to the top 10 components. To assign domains, self-organizing maps (SOM) were used for low-rank quantization of query embeddings. In analysis of the MERFISH dataset, SOMs of size 1 x k were fit across a range of 2 to 12; the best model was determined using the elbow method heuristic to evaluate quantization error. Similarly, domains were determined for the cardiomyocytes spatial transcriptomics data by fitting the vehicle and treatment samples separately, for k across a range of 2 to 8. The elbow method heuristic determined an optimal k of 6; subsequently a k of 4 was used for further analysis for ease of interpretation.

### RNAflux: Visualizing spatial embeddings

The top 3 principal components of the RNAflux embeddings are transformed to map to red, green and blue values respectively. Embeddings are first quantile normalized and scaled to a minimum of 0.1 and 0.9 to avoid mapping extreme quantiles to white and black. These values are then used for red, green, and blue color channels. To map the downsampled grid back to the original data units, linear interpolation was used to rescale the computed color values and fill the space between the uniform grid points.

### RNAflux: Enrichment of organelle-specific transcriptomes

The enrichment score for each pixel is calculated by first taking the weighted sum of its RNAflux embedding and published compartment log fold-change values as implemented by the decoupler tool^70^. Scores for pixels within a given cell are normalized against a null distribution constructed via random permutations of the input embeddings, to produce z-scaled enrichment scores. Fluxmap domain enrichment scores are simply obtained by taking the mean score of all pixels within the boundary of each domain.

### RNAForest: model selection and training

We evaluated 4 base models for the multilabel classifier including random forests (RF), support vector machines (SVM), feed-forward fully-connected neural networks (NN), and convolutional neural networks (CNN). While all other models use the 13 spatial features for input (**Supp. Table 1**), the CNN takes 64x64 image representations of each sample as input. Each multilabel classifier consists of 5 binary classifiers with the same base model. We used the labeled 10,000 simulated samples for training, stratifying 80% of the simulated data for training and holding out the remaining 20% for testing. To select the best hyperparameters for each multilabel classifier, we sampled from a fixed hyperparameter space with the Tree-structured Parzen Estimator algorithm, and evaluated performance with 5-fold cross validation (**Supp. Table 3**). We retrained the final model (random forest base model) on all training data with the best performing set of hyperparameters (**Supp. Fig. 2E**).

### RNAforest: Image rasterization of molecules and segmentation masks for CNN

To generate an image for a given sample, point coordinates, the cell segmentation mask and nuclear segmentation mask are used. The area of the cell is tiled as a 64 x 64 grid, where each bin corresponds to a pixel in the final image. Values are stored in a single channel to render a grayscale image. Pixels inside the cell are encoded as 20, inside the nucleus encoded as 40. Bins with molecules are encoded as (40 + 20 x n) where n is the number of molecules. Finally values are divided by 255 and capped to be between 0 and 1.

### RNAforest: Simulating training data

We trained a multilabel classifier to assign each gene in every cell labels from five categories: (i) nuclear (contained in the volume of the nucleus), (ii) cytoplasmic (diffuse throughout the cytoplasm), (iii) nuclear edge (near the inner/outer nuclear membrane), (iv) cell edge (near the cell membrane), and (v) none (complete spatial randomness). These categories are a consolidation of those observed in several high-throughput smFISH imaging experiments in HeLa cells ^40–43^. We used the FISH-quant simulation framework to generate realistic ground-truth images using empirically derived parameters from the mentioned high-throughput smFISH imaging experiments in HeLa cells ^42^. In total, we simulate 2,000 samples per class for a total of 10,000 training samples.

1. **Cell shape**: Cell morphology varies widely across cell types and for classifier generalizability, it is important to include many different morphologies in the training set. We use a catalog of cell shapes for over 300 cells from smFISH images in HeLa cells that captures nucleus and cell membrane shape ^42^. Cell shapes were obtained by cell segmentation with CellMask and nuclear segmentation was obtained from DAPI staining.
2. **mRNA abundance**: We simulated mRNA abundance at three different expression levels (40, 100, and 200 mRNA per average sized cell) with a Poisson noise term. Consequently, total mRNA abundance per cell was between 5 and 300 transcripts.
3. **Localization pattern**: We focused on 5 possible 2D localization patterns, including cell edge, cytoplasmic, none, nuclear, and nuclear edge. Each pattern was further evaluated at 3 different degrees - weak, moderate, and strong. Moderate corresponds to a pattern typically observed in a cell, whereas weak is close to spatially random. These 5 classes aim to capture biologically relevant behavior generalizable to most cell types; there is room for additional classes describing other biologically relevant localization patterns so long as they can be accurately modeled.

### RNAforest: Manual annotation of validation data

Using 3 individual annotators, we annotated the same 600 samples across both datasets, keeping samples with 2 or more annotator agreements as true annotations, resulting in 165 annotated seqFISH+ samples and 238 annotated MERFISH samples (403 total). We used Cohen’s kappa coefficient^71^ to calculate agreement between pairs of annotators for each label yielding an overall coefficient of 0.602. We found that pairwise agreement between annotators across labels was fairly consistent ranging between 0.588 and 0.628, while label-specific agreement varied more, ranging between 0.45 and 0.72 (**Supp. Table 4**).

### RNAforest: Functional enrichment of gene pattern distributions

For enrichment of compartment-specific expression from Xia et al 2019^46^, scores are calculated by taking the weighted sum of gene pattern frequencies and published compartment log fold-change values (**Supp. Fig. 3**). The Benjamini-Hochberg correction was used to correct p-values for multiple hypothesis testing.

For the seqFISH+ dataset, we performed single-sample Gene Set Enrichment Analysis^72, 73^ on gene pattern frequencies to compute enrichment scores (**Fig. 3I**). ssGSEA was performed with the GSEApy Python package and the “GO_Cellular_Component_2021” gene set library curated by Enrichr^74^. Gene sets with a minimum size of 50 and a maximum size of 500 were analyzed.

### Colocation quotient for RNA colocalization analysis

Pairwise colocalization of genes was determined for each compartment of every cell separately. In this case, each cell was divided into compartments, cytoplasm and nucleus. The colocation quotient (CLQ) was calculated for every pair of genes 𝐴 and 𝐵. The CLQ is defined as an odds ratio of the observed to expected proportion of 𝐵 transcripts among neighbors of 𝐴 for a fixed radius r; it is formulated as:

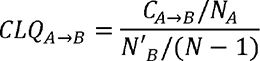

Here 𝐶_𝐴_*_→_*_𝐵_ denotes the number of 𝐴 transcripts of which 𝐵 transcripts are considered a neighbor. 𝑁_𝐴_ denotes the total number of *A* transcripts, while N’B stands for the total number of 𝐵 transcripts. In the case that 𝐴 = 𝐵, N’B equals the total number of 𝐵 transcripts minus one. 𝑁 denotes the total number of transcripts in the cell. Following statistical recommendations from the original formulation of the colocation quotient (CLQ), genes with fewer than 10 transcripts were not considered to reduce sparsity and improve testing power^51^.

### Tensor decomposition for compartment-specific colocalization

For tensor decomposition, we employed non-negative parallel factor analysis as implemented in Tensorly^31^, which seeks to represent our dataset tensor **X** in a lower dimensional space of **R** signatures by decomposing **X** as the sum of **R** rank-one 3-way tensors. Each of these tensors is described as the outer product of 3 vectors, 𝑥^𝑟^, 𝑦^𝑟^ and 𝑧^𝑟^ . The collection of vectors across **R** signatures we denote as 𝑥^𝑟^ (compartment loadings), 𝑦^𝑟^ (cell loadings) and 𝑧^𝑟^ (gene pair loadings) respectively. We find the optimal rank-**R** decomposition of **X** by minimizing reconstruction error as a function of the number of signatures **R** and use the elbow function heuristic to choose the best-fit across the range of 2-12 factors.

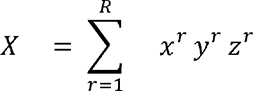

### MERFISH of U2-OS cells

#### MERFISH sample preparation

MERFISH measurements of 130 genes with five non-targeting blank controls was done as previously described, using the published encoding^44^ and readout probes^75^. Briefly, U2-OS cells were cultured on 40 mm #1.5 coverslips that are silanized and poly-L-lysine coated^44^ and subsequently fixed in 4% (vol/vol) paraformaldehyde in 1x PBS for 15 minutes at room temperature. Cells were then permeabilized in 0.5% Triton X-100 for 10 minutes at room temperature and washed in 1x PBS containing Murine RNase Inhibitor (NEB M0314S). Cells were preincubated with hybridization wash buffer (30% (vol/vol) formamide in 2x SSC) for ten minutes at room temperature with gentle shaking. After preincubation, the coverslip was moved to a fresh 60 mm petri dish and residual hybridization wash buffer was removed with a Kimwipe lab tissue. In the new dish, 50 uL of encoding probe hybridization buffer (2X SSC, 30% (vol/vol) formamide, 10% (wt/vol) dextran sulfate, 1 mg ml^-1^ yeast tRNA, and a total concentration of 5 uM encoding probes and 1 μM of anchor probe: a 15-nt sequence of alternating dT and thymidine-locked nucleic acid (dT+) with a 5′-acrydite modification (Integrated DNA Technologies). The sample was placed in a humidified 37C oven for 36 to 48 hours then washed with 30% (vol/vol) formamide in 2X SSC for 20 minutes at 37C, 20 minutes at room temperature. Samples were post-fixed with 4% (vol/vol) paraformaldehyde in 2X SSC and washed with 2X SSC with murine RNase inhibitor for five minutes. The samples were finally stained with a Alexa 488-conjugated anchor probe-readout oligo (Integrated DNA Technologies) and DAPI solution at 1 μg/ml.

#### MERFISH imaging

MERFISH measurements were conducted on a home-built system as described in Huang et al. 2021^75^.

#### MERFISH spot detection

Individual RNA molecules were decoded in MERFISH images using MERlin v0.1.6^76^. Images were aligned across hybridization rounds by maximizing phase cross-correlation on the fiducial bead channel to adjust for drift in the position of the stage from round to round. Background was reduced by applying a high-pass filter and decoding was then performed per-pixel. For each pixel, a vector was constructed of the 16 brightness values from each of the 16 rounds of imaging. These vectors were then L2 normalized and their euclidean distances to each of the L2 normalized barcodes from MERFISH codebook was calculated. Pixels were assigned to the gene whose barcode they were closest to, unless the closest distance was greater than 0.512, in which case the pixel was not assigned a gene. Adjacent pixels assigned to the same gene were combined into a single RNA molecule. Molecules were filtered to remove potential false positives by comparing the mean brightness, pixel size, and distance to the closest barcode of molecules assigned to blank barcodes to those assigned to genes to achieve an estimated misidentification rate of 5%. The exact position of each molecule was calculated as the median position of all pixels consisting of the molecule.

#### MERFISH image segmentation

Cellpose v1.0.2^63^ was used to perform image segmentation to determine the boundaries of cells and nuclei. The nuclei boundaries were determined by running Cellpose with the ‘nuclei’ model using default parameters on the DAPI stain channel of the pre-hybridization images. Cytoplasm boundaries were segmented with the ‘cyto’ model and default parameters using the polyT stain channel. RNA molecules identified by MERlin were assigned to cells and nuclei by applying these segmentation masks to the positions of the molecules.

### iPSC Cardiac Differentiation and Doxorubicin Treatment

Matrigel (Corning, cat # 354277) coated plates were used to culture iPSCs with mTESR Plus human iPSC medium (StemCell Technologies, cat # 100-0276) in a humidified incubator at 37°C with 5% CO_2_. iPSCs were dissociated with Gentle Cell Dissociation Reagent (StemCell Technologies, cat # 100-0485) and passaged with mTESR Plus medium and 10uM ROCK inhibitor (Tocris, cat #1254) at a ratio of 1:12. mTESR plus medium was replaced every other day until the cells reached 80% confluency for maintenance and replating, or 90% confluency for cardiac differentiation utilizing a chemically defined protocol^77^. On day 0 of cardiac differentiation, cells were treated with 6μM CHIR99021 (Selleck Chem, cat # S1263) in RPMI 1640 media (Gibco, cat # 11875) and B27 minus insulin supplement (Thermo Fisher, cat # A1895601). On day 2, CHIR was removed, and cells were cultured with RPMI 1640 media and B27 minus insulin supplement (Thermo Fisher, cat # A18956). On day 3, media was replaced with RPMI media containing B27 minus insulin supplement and 5 μM Wnt-C59 (Cellagen Technologies, cat # C7641-2s). On days 5, 7, and 9, media was replaced with RPMI media containing B27 insulin supplement (Thermo Fisher, cat # 17504). On days 11 and 13, media was replaced with RPMI 1640 media without glucose (Thermo Fisher, cat # 11879020) containing B27 insulin supplement for purification of cardiomyocytes. From days 15 onward, the cells were cultured in RPMI 1640 media containing B27 supplement which was changed every other day until the cells reached day 30 for replating. For replating, cells were incubated in 10X TrypLE (Thermo Fisher, cat # A1217701) for 12 minutes at 37 C, neutralized with equal volumes of RPMI 1640 media containing B27 supplement with 20% FBS (Gibco, cat # 26140-079), gently dissociated by pipetting, then spun down and resuspended for replating in RPMI 1640 media containing B27 supplement with 20% FBS. The next day, the cell media was replaced with RPMI 1640 media containing B27 supplement which was replaced with fresh media every other day. On day 48 the cells were replated onto chamber slides (Ibidi, cat # 80826) as described above and recovered for 10 days before doxorubicin treatments began (MedChemExpress, cat # HY-15142). On day 60, doxorubicin treatments concluded, and the cells underwent methanol fixation.

### Molecular Cartography

#### Cultured cell processing

After Doxorubicin treatment, cardiomyocytes were washed with PBS (1x) twice and fixed in Methanol (−20°C) for 10 min. After fixation, Methanol was aspirated and cells were dried and stored at −80°C until use. The samples were used for Molecular Cartography^TM^ (100-plex combinatorial single molecule fluorescence in-situ hybridization) according to the manufacturer’s instructions *Day 1: Molecular Preparation Protocol* for cells, starting with the addition of buffer DST1 followed by cell priming and hybridization. Briefly, cells were primed for 30 minutes at 37°C followed by overnight hybridization of all probes specific for the target genes (see below for probe design details and target list). Samples were washed the next day to remove excess probes and fluorescently tagged in a two-step color development process. Regions of interest were imaged as described below and fluorescent signals removed during decolorization. Color development, imaging and decolorization were repeated for multiple cycles to build a unique combinatorial code for every target gene that was derived from raw images as described below.

#### Probe Design

The probes for 100 genes were designed using Resolve’s proprietary design algorithm. Briefly, the probe-design was performed at the gene-level. For every targeted gene, all full-length protein coding transcript sequences from the ENSEMBL database were used as design targets if the isoform had the GENCODE annotation tag ‘basic’^78, 79^. To speed up the process, the calculation of computationally expensive parts, especially the off-target searches, the selection of probe sequences was not performed randomly, but limited to sequences with high success rates. To filter highly repetitive regions, the abundance of k-mers was obtained from the background transcriptome using Jellyfish^80^. Every target sequence was scanned once for all k-mers, and those regions with rare k-mers were preferred as seeds for full probe design. A probe candidate was generated by extending a seed sequence until a certain target stability was reached. A set of simple rules was applied to discard sequences that were found experimentally to cause problems. After these fast screens, the remaining probe candidates were mapped to the background transcriptome using ThermonucleotideBLAST^81^ and probes with stable off-target hits were discarded. Specific probes were then scored based on the number of on-target matches (isoforms), which were weighted by their associated APPRIS level^82^, favoring principal isoforms over others. A bonus was added if the binding-site was inside the protein-coding region. From the pool of accepted probes, the final set was composed by picking the highest scoring probes. Probes with catalog numbers can be found in **Supp. Table 2**.

#### Imaging

Samples were imaged on a Zeiss Celldiscoverer 7, using the 50x Plan Apochromat water immersion objective with an NA of 1.2 and the 0.5x magnification changer, resulting in a 25x final magnification. Standard CD7 LED excitation light source, filters, and dichroic mirrors were used together with customized emission filters optimized for detecting specific signals. Excitation time per image was 1000 ms for each channel (DAPI was 20 ms). A z-stack was taken at each region with a distance per z-slice according to the Nyquist-Shannon sampling theorem. The custom CD7 CMOS camera (Zeiss Axiocam Mono 712, 3.45 µm pixel size) was used. For each region, a z-stack per fluorescent color (two colors) was imaged per imaging round. A total of 8 imaging rounds were done for each position, resulting in 16 z-stacks per region. The completely automated imaging process per round was realized by a custom python script using the scripting API of the Zeiss ZEN software (Open application development).

#### Image Processing and Spot Segmentation

As a first step all images were corrected for background fluorescence. A target value for the allowed number of maxima was determined based upon the area of the slice in µm² multiplied by the factor 0.5. This factor was empirically optimized. The brightest maxima per plane were determined, based upon an empirically optimized threshold. The number and location of the respective maxima was stored. This procedure was done for every image slice independently. Maxima that did not have a neighboring maximum in an adjacent slice (called z-group) were excluded. The resulting maxima list was further filtered in an iterative loop by adjusting the allowed thresholds for (Babs-Bback) and (Bperi-Bback) to reach a feature target value (Babs: absolute brightness, Bback: local background, Bperi: background of periphery within 1 pixel). This feature target values were based upon the volume of the 3D-image. Only maxima still in a zgroup of at least 2 after filtering were passing the filter step. Each z-group was counted as one hit. The members of the z-groups with the highest absolute brightness were used as features and written to a file. They resemble a 3D-point cloud. To align the raw data images from different imaging rounds, images had to be registered. To do so the extracted feature point clouds were used to find the transformation matrices. For this purpose, an iterative closest point cloud algorithm was used to minimize the error between two point-clouds. The point clouds of each round were aligned to the point cloud of round one (reference point cloud). The corresponding point clouds were stored for downstream processes. Based upon the transformation matrices the corresponding images were processed by a rigid transformation using trilinear interpolation. The aligned images were used to create a profile for each pixel consisting of 16 values (16 images from two color channels in 8 imaging rounds). The pixel profiles were filtered for variance from zero normalized by total brightness of all pixels in the profile. Matched pixel profiles with the highest score were assigned as an ID to the pixel. Pixels with neighbors having the same ID were grouped. The pixel groups were filtered by group size, number of direct adjacent pixels in group, number of dimensions with size of two pixels. The local 3D-maxima of the groups were determined as potential final transcript locations. Maxima were filtered by the number of maxima in the raw data images where a maximum was expected. Remaining maxima were further evaluated by the fit to the corresponding code. The remaining maxima were written to the results file and considered to resemble transcripts of the corresponding gene. The ratio of signals matching to codes used in the experiment and signals matching to codes not used in the experiment were used as estimation for specificity (false positives). The algorithms for spot segmentation were written in Java and are based on the ImageJ library functionalities. Only the iterative closest point algorithm is written in C++ based on the libpointmatcher library (https://github.com/ethz-asl/libpointmatcher).

#### Image segmentation

Cellpose v1.0.2^63^ was used to perform image segmentation to determine the boundaries of nuclei. The nuclei boundaries were determined by running Cellpose with the ‘nuclei’ model using default parameters on the DAPI stain channel of the pre-hybridization images. Cytoplasm boundaries were determined with ClusterMap^24^ using spot coordinates.

## Data Availability

Preprocessed and raw datasets have been deposited at https://doi.org/10.6084/m9.figshare.c.6564043.v1 and are accessible through the Bento Python package. These include the seqFISH+^14^ and the generated MERFISH datasets. Raw MERFISH data is available upon request.

## Code Availability

The source code for Bento is available on the GitHub repository: https://github.com/ckmah/bento-tools. Analysis code for generating figures can be found at: https://github.com/ckmah/bento-manuscript.

Documentation for Bento can be found here: http://bento-tools.readthedocs.io/.

## Acknowledgements

C.K.M. is supported by the National Science Foundation Graduate Research Fellowship under Grant No. (DGE-2038238). N.A. was partially supported by NIH Training Grant T32 GM008666. This work was partially supported by National Institutes of Health grants NS103172, MH107367, AI132122, AI123202, AG069098, HG004659, and HG009889 to G.W.Y. G.W.Y. is also supported by an Allen Distinguished Investigator Award, a Paul G. Allen Frontiers Group advised grant of the Paul G. Allen Family Foundation. A.J.C. and E.L. acknowledge support from the Chan Zuckerberg Initiative (CZF2019-002448) and the Knut and Alice Wallenberg Foundation (KAW 2021.0346) to E.L. We thank members of the Yeo lab, Carter lab, Michelle Franc Ragsac, Erick Armingol, and Nate Lewis for helpful discussions and feedback on the manuscript.

## Author Contributions

C.K.M, N.A., and G.W.Y. conceptualized the project. C.K.M. and N.A. co-developed the software. C.K.M. and D.L. trained the classification model for subcellular localization. C.K.M., N.A., and D.L. manually annotated data for benchmarking model performance. C.K.M., N.A. and G.P. performed data preprocessing and analysis. A.M., C.K., Y.H., and Q.Z. generated the MERFISH experiment. N.L. designed the gene panel and cultured the cardiomyocytes. A.C. and E.L. aided multimodal spatial analyses. C.K.M., N.A., H.C., and G.W.Y. wrote the manuscript. H.C. and G.W.Y. supervised the project.

## Supplementary Figures

**Fig S1.**
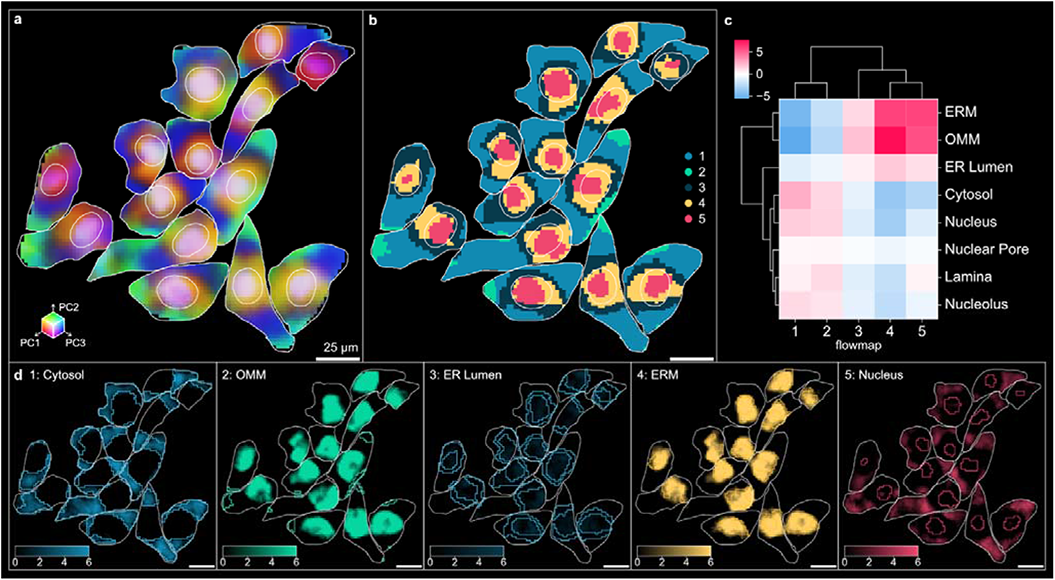
RNAflux characterizes subcellular regions in seqFISH+ 3T3 cells. **A)** The first three principal components of the RNAflux embedding are visualized for seqFISH+ cells coloring RGB values by PC1, PC2, and PC3 values respectively for each pixel. **B)** Fluxmap domains are computed from each RNAflux embedding to create semantic segmentation masks of each subcellular domain. **C)** Mean enrichment score of pixels contained within each fluxmap domain (**Methods**). **D)** Relative enrichment of transcripts enriched for organelle-specific expression, as processed for the MERFISH U2-OS dataset, are highlighted along with the SOM segmentation boundary for the fluxmap domain expected to be enriched based on its location in the cell. Stronger intensity denotes positive enrichment while black denotes no enrichment.

**Fig S2.**
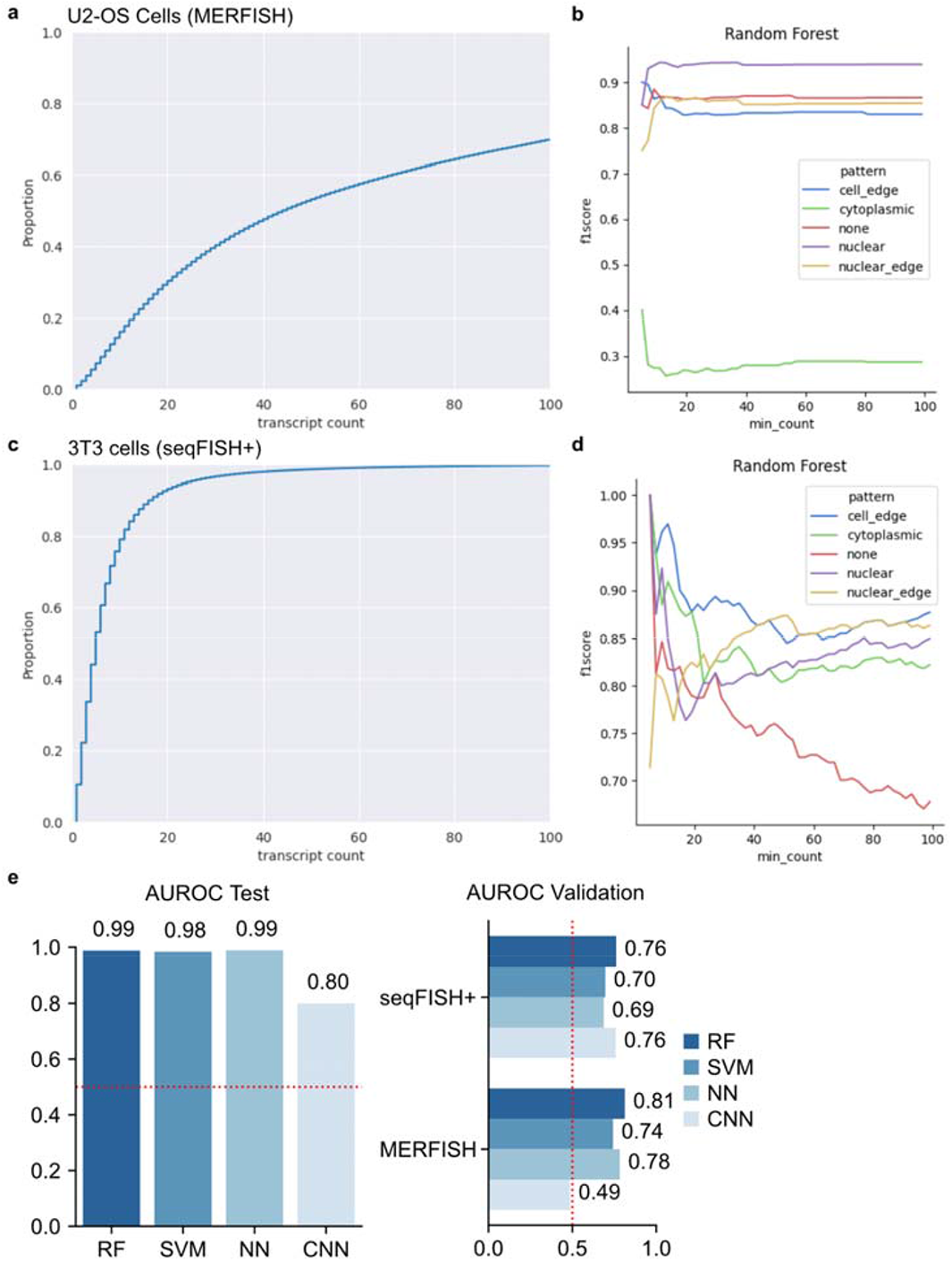
RNAforest performance evaluation. **A)** Cumulative distribution of sample molecule copy number in U2-OS cells MERFISH dataset. **B)** Validation F1-score of each binary classifier in RNAforest as a function of sample molecule copy number for MERFISH dataset. **C)** Cumulative distribution of sample molecule copy number in 3T3 cells seqFISH+ dataset. **D)** Validation F1-score of each binary classifier in RNAforest as a function of sample molecule copy number for seqFISH+ dataset. **E)** Benchmarking performance of the 4 base models (RF - random forest, SVM - support vector machine, NN - fully connected neural network, CNN - convolutional neural network), showing AUROC in test and validation data.

**Fig S3.**
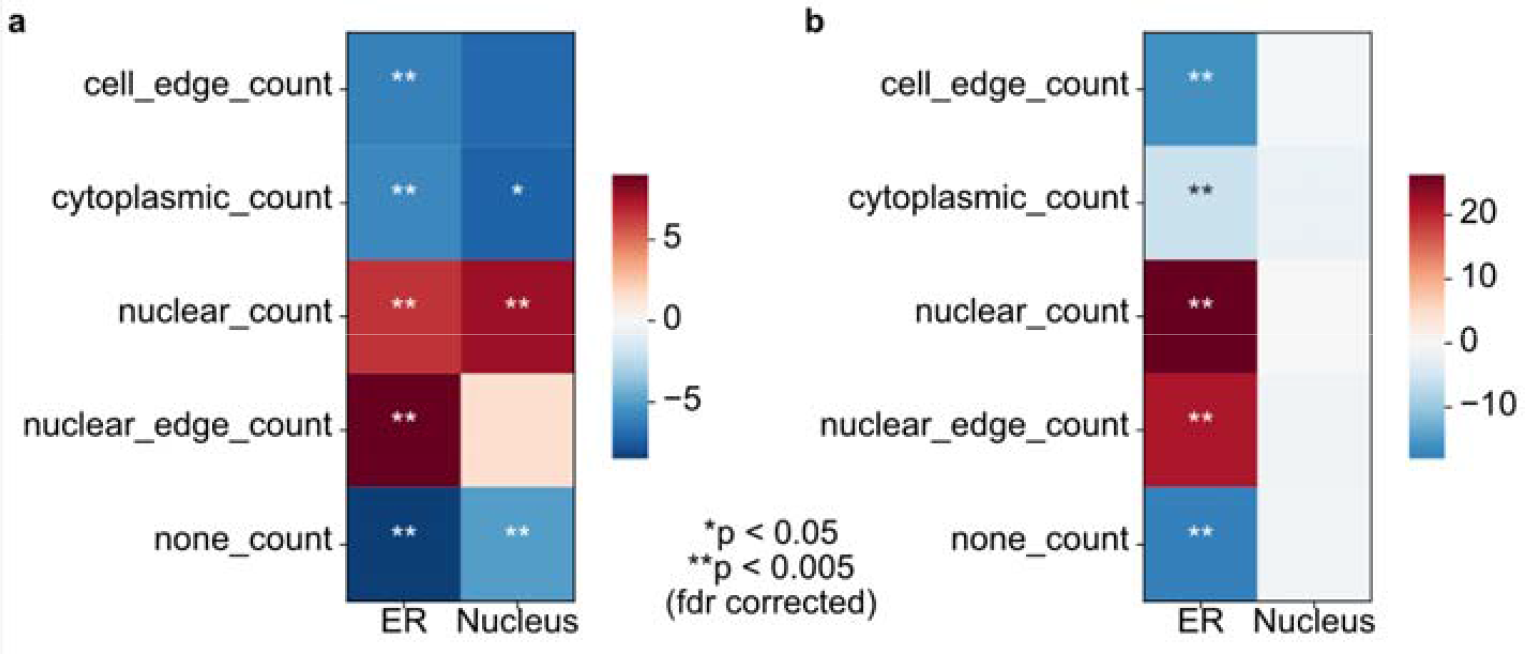
Enrichment of compartment-specific expression for RNAforest gene pattern frequencies. Compartment-specific enrichment of endoplasmic reticulum (ER) and nucleus gene expression – from Xia et al 2019^46^ – relative to RNAforest gene pattern frequencies in the **A)** MERFISH dataset and **B)** seqFISH+ dataset.

**Fig S4.**
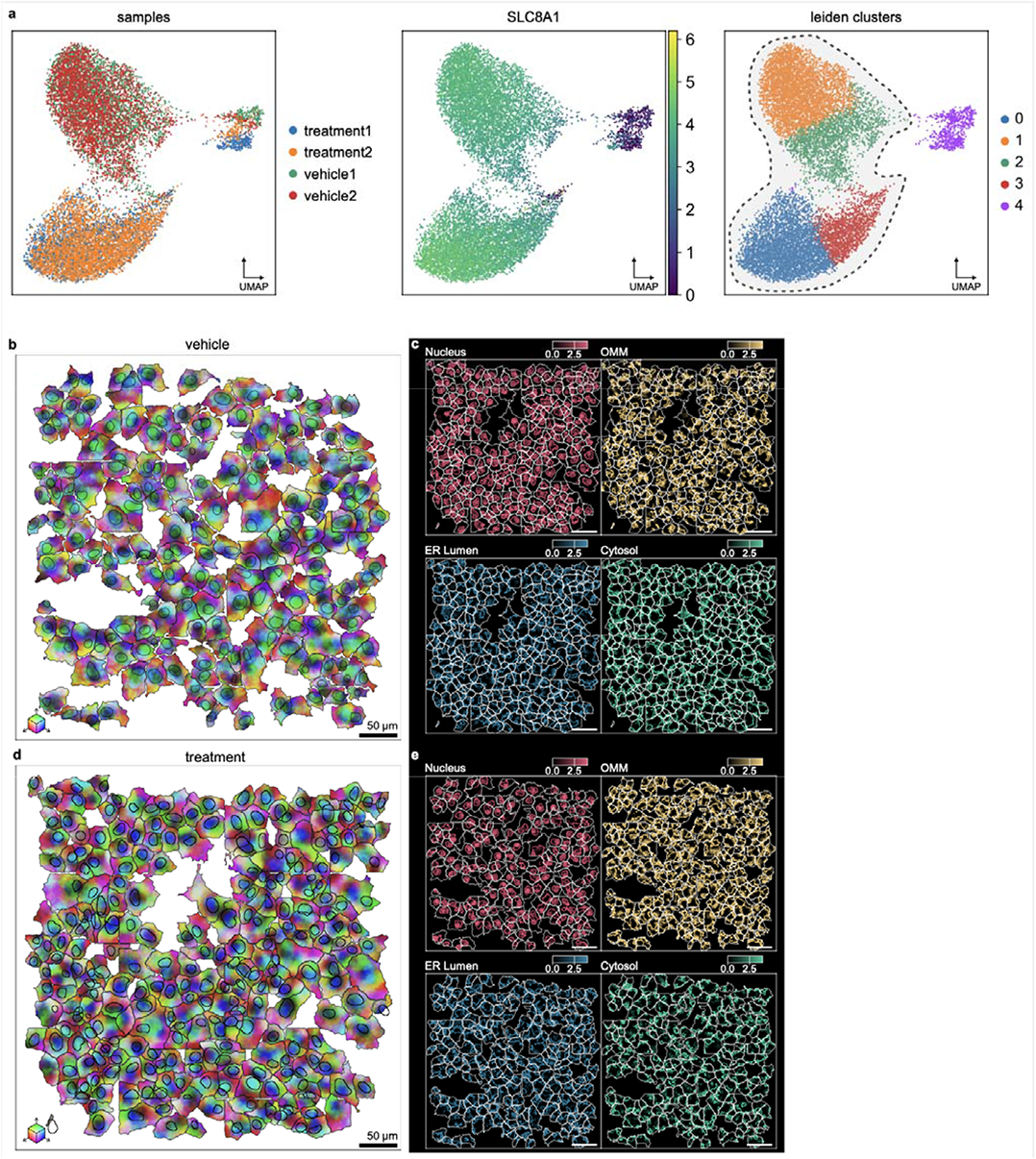
Filtering and RNAflux analysis of DOX treated cardiomyocytes. **A)** Left: UMAP of all 4 cardiomyocyte samples, colors denote different samples. Center: Cells are colored by log-scaled SLC8A1 RNA expression. Right: Leiden clustering identifies 5 clusters, separating low expression SLC8A1 into cluster 4. Representative crop of **B)** vehicle and **D)** treatment samples, colored by the first 3 principal components of its RNAflux embedding. Relative enrichment of transcripts enriched for organelle-specific expression in **C)** vehicle and **E)** treatment samples. Red, yellow, blue and green enrichment correspond to nuclear, OMM, ER lumen, and cytosol genesets respectively.

## References

1. Thul, P. J. et al. A subcellular map of the human proteome. Science 356, (2017).

2. Laurila, K. & Vihinen, M. Prediction of disease-related mutations affecting protein localization. BMC

3. Park, S. et al. Protein localization as a principal feature of the etiology and comorbidity of genetic diseases. Mol. Syst. Biol. 7, 494 (2011).

4. Chin, A. & Lécuyer, E. RNA localization: Making its way to the center stage. Biochim. Biophys. Acta Gen. Subj. 1861, 2956–2970 (2017).

5. Bovaird, S., Patel, D., Padilla, J.-C. A. & Lécuyer, E. Biological functions, regulatory mechanisms, and disease relevance of RNA localization pathways. FEBS Lett. 592, 2948–2972 (2018).

6. Das, S., Singer, R. H. & Yoon, Y. J. The travels of mRNAs in neurons: do they know where they are going? Curr. Opin. Neurobiol. 57, 110–116 (2019).

7. Sahoo, P. K., Smith, D. S., Perrone-Bizzozero, N. & Twiss, J. L. Axonal mRNA transport and translation at a glance. J. Cell Sci. 131, (2018).

8. von Kügelgen, N. & Chekulaeva, M. Conservation of a core neurite transcriptome across neuronal types and species. Wiley Interdiscip. Rev. RNA e1590 (2020).

9. Culver, B. P. et al. Huntington’s Disease Protein Huntingtin Associates with its own mRNA. J Huntingtons Dis 5, 39–51 (2016).

10. Romo, L., Mohn, E. S. & Aronin, N. A Fresh Look at Huntingtin mRNA Processing in Huntington’s Disease. J Huntingtons Dis 7, 101–108 (2018).

11. White, J. A., 2nd et al. Huntingtin differentially regulates the axonal transport of a sub-set of Rab-containing vesicles in vivo. Hum. Mol. Genet. 24, 7182–7195 (2015).

12. Fernandopulle, M. S., Lippincott-Schwartz, J. & Ward, M. E. RNA transport and local translation in neurodevelopmental and neurodegenerative disease. Nat. Neurosci. (2021) doi:10.1038/s41593-020-00785-2.

13. Chen, K. H., Boettiger, A. N., Moffitt, J. R., Wang, S. & Zhuang, X. RNA imaging. Spatially resolved, highly multiplexed RNA profiling in single cells. Science 348, aaa6090 (2015).

14. Eng, C.-H. L. et al. Transcriptome-scale super-resolved imaging in tissues by RNA seqFISH. Nature 568, 235–239 (2019).

15. Gyllborg, D. et al. Hybridization-based in situ sequencing (HybISS) for spatially resolved transcriptomics in human and mouse brain tissue. Nucleic Acids Res. (2020) doi:10.1093/nar/gkaa792.

16. Alon, S. et al. Expansion Sequencing: Spatially Precise In Situ Transcriptomics in Intact Biological Systems. Cold Spring Harbor Laboratory 2020.05.13.094268 (2020) doi:10.1101/2020.05.13.094268.

17. Dries, R. et al. Giotto: a toolbox for integrative analysis and visualization of spatial expression data. Genome Biol. 22, 78 (2021).

18. Pham, D. et al. stLearn: integrating spatial location, tissue morphology and gene expression to find cell types, cell-cell interactions and spatial trajectories within undissociated tissues. bioRxiv 2020.05.31.125658 (2020) doi:10.1101/2020.05.31.125658.

19. Butler, A., Hoffman, P., Smibert, P., Papalexi, E. & Satija, R. Integrating single-cell transcriptomic data across different conditions, technologies, and species. Nat. Biotechnol. 36, 411–420 (2018).

20. Wolf, F. A., Angerer, P. & Theis, F. J. SCANPY: large-scale single-cell gene expression data analysis. Genome Biol. 19, 15–15 (2018).

21. Imbert, A. et al. FISH-quant v2: a scalable and modular tool for smFISH image analysis. RNA 28, 786– 795 (2022).

22. Walter, F. C., Stegle, O. & Velten, B. FISHFactor: A probabilistic factor model for spatial transcriptomics data with subcellular resolution. bioRxiv (2021) doi:10.1101/2021.11.04.467354.

23. Imbert, A. et al. FISH-quant v2: a scalable and modular analysis tool for smFISH image analysis. bioRxiv 2021.07.20.453024 (2021) doi:10.1101/2021.07.20.453024.

24. He, Y. et al. ClusterMap for multi-scale clustering analysis of spatial gene expression. Nat. Commun. 12, 5909 (2021).

25. Petukhov, V. et al. Cell segmentation in imaging-based spatial transcriptomics. Nat. Biotechnol. 40, 345–354 (2022).

26. Spitzer, H., Berry, S., Donoghoe, M., Pelkmans, L. & Theis, F. J. Learning consistent subcellular landmarks to quantify changes in multiplexed protein maps. bioRxiv 2022.05.07.490900 (2022) doi:10.1101/2022.05.07.490900.

27. Liu, C. C. et al. Robust phenotyping of highly multiplexed tissue imaging data using pixel-level clustering. bioRxiv 2022.08.16.504171 (2022) doi:10.1101/2022.08.16.504171.

28. Jordahl, K. et al. geopandas/geopandas: v0.9.0. (2021). doi:10.5281/zenodo.4569086.

29. Gillies, S., Ward, B. & Petersen, A. S. Rasterio: Geospatial raster I/O for Python programmers. *URL* https://github.com/mapbox/rasterio.

30. Virtanen, P. et al. SciPy 1.0: fundamental algorithms for scientific computing in Python. Nat. Methods 17, 261–272 (2020).

31. Kossaifi, J., Panagakis, Y., Anandkumar, A. & Pantic, M. TensorLy: Tensor Learning in Python. J. Mach. Learn. Res. 20, 1–6 (2019).

32. Palla, G. et al. Squidpy: a scalable framework for spatial omics analysis. Nat. Methods (2022) doi:10.1038/s41592-021-01358-2.

33. He, S. et al. High-plex multiomic analysis in FFPE at subcellular level by spatial molecular imaging. bioRxiv (2021) doi:10.1101/2021.11.03.467020.

34. Lee, J. H. et al. Fluorescent in situ sequencing (FISSEQ) of RNA for gene expression profiling in intact cells and tissues. Nat. Protoc. 10, 442–458 (2015).

35. Hu, S. et al. Dynamic control of metabolic zonation and liver repair by endothelial cell Wnt2 and Wnt9b revealed by single cell spatial transcriptomics using Molecular Cartography. bioRxiv 2022.03.18.484868 (2022) doi:10.1101/2022.03.18.484868.

36. Virshup, I., Rybakov, S., Theis, F. J., Angerer, P. & Alexander Wolf, F. anndata: Annotated data. bioRxiv 2021.12.16.473007 (2021) doi:10.1101/2021.12.16.473007.

37. Volkova, M. & Russell, R., 3rd. Anthracycline cardiotoxicity: prevalence, pathogenesis and treatment. Curr. Cardiol. Rev. 7, 214–220 (2011).

38. Fazal, F. M. et al. Atlas of Subcellular RNA Localization Revealed by APEX-Seq. Cell 178, 473– 490.e26 (2019).

39. Viola, P. & Jones, M. Rapid object detection using a boosted cascade of simple features. in *Proceedings of the 2001 IEEE Computer Society Conference on Computer Vision and Pattern Recognition*. CVPR 2001 vol. 1 I–I (2001).

40. Battich, N., Stoeger, T. & Pelkmans, L. Image-based transcriptomics in thousands of single human cells at single-molecule resolution. Nat. Methods 10, 1127–1133 (2013).

41. Stoeger, T., Battich, N., Herrmann, M. D., Yakimovich, Y. & Pelkmans, L. Computer vision for image-based transcriptomics. Methods 85, 44–53 (2015).

42. Samacoits, A. et al. A computational framework to study sub-cellular RNA localization. Nat. Commun. 9, 4584 (2018).

43. Chouaib, R. et al. A Dual Protein-mRNA Localization Screen Reveals Compartmentalized Translation and Widespread Co-translational RNA Targeting. Dev. Cell 54, 773–791.e5 (2020).

44. Moffitt, J. R. et al. High-throughput single-cell gene-expression profiling with multiplexed error-robust fluorescence in situ hybridization. Proc. Natl. Acad. Sci. U. S. A. 113, 11046–11051 (2016).

45. Kumar, A. et al. Intracellular Spatial Transcriptomic Analysis Toolkit (InSTAnT). Research Square (2023) doi:10.21203/rs.3.rs-2481749/v1.

46. Xia, C., Fan, J., Emanuel, G., Hao, J. & Zhuang, X. Spatial transcriptome profiling by MERFISH reveals subcellular RNA compartmentalization and cell cycle-dependent gene expression. Proc. Natl. Acad. Sci. U. S. A. 116, 19490–19499 (2019).

47. Gene Ontology Consortium. The Gene Ontology resource: enriching a GOld mine. Nucleic Acids Res. 49, D325–D334 (2021).

48. Ripley, B. D. The second-order analysis of stationary point processes. J. Appl. Probab. 13, 255–266 (1976).

49. Tiefelsdorf, M. Modelling Spatial Processes: The Identification and Analysis of Spatial Relationships in Regression Residuals by Means of Moran’s I. (Springer, 2006).

50. Cliff, A. D. & Ord, J. K. Spatial Processes: Models & Applications. (Pion, 1981).

51. Leslie, T. F. &Kronenfeld, B. J. The colocation quotient: A new measure of spatial association between categorical subsets of points. •同区位商 : 点集分•子集•空•••性的新度量•准: The colocation quotient. Geogr. Anal. 43, 306–326 (2011).

52. Shashua, A. & Hazan, T. Non-negative tensor factorization with applications to statistics and computer vision. in Proceedings of the 22nd international conference on Machine learning 792–799 (Association for Computing Machinery, 2005).

53. Young, R. C., Ozols, R. F. & Myers, C. E. The anthracycline antineoplastic drugs. N. Engl. J. Med. 305, 139–153 (1981).

54. Kalyanaraman, B. Teaching the basics of the mechanism of doxorubicin-induced cardiotoxicity: Have we been barking up the wrong tree? Redox Biol 29, 101394 (2020).

55. Sheibani, M. et al. Doxorubicin-Induced Cardiotoxicity: An Overview on Pre-clinical Therapeutic Approaches. Cardiovasc. Toxicol. 22, 292–310 (2022).

56. Yu, J. et al. Recent progress in doxorubicin-induced cardiotoxicity and protective potential of natural products. Phytomedicine 40, 125–139 (2018).

57. Rahman, A. M., Yusuf, S. W. & Ewer, M. S. Anthracycline-induced cardiotoxicity and the cardiac-sparing effect of liposomal formulation. Int. J. Nanomedicine 2, 567–583 (2007).

58. Xu, M. F., Tang, P. L., Qian, Z. M. & Ashraf, M. Effects by doxorubicin on the myocardium are mediated by oxygen free radicals. Life Sci. 68, 889–901 (2001).

59. Šimůnek, T., et al. Anthracycline-induced cardiotoxicity: Overview of studies examining the roles of oxidative stress and free cellular iron. Pharmacol. Rep. 61, 154–171 (2009).

60. Xiong, C. et al. Protective effect of berberine on acute cardiomyopathy associated with doxorubicin treatment. Oncol. Lett. 15, 5721–5729 (2018).

61. Asensio-López, M. C., Soler, F., Pascual-Figal, D., Fernández-Belda, F. & Lax, A. Doxorubicin-induced oxidative stress: The protective effect of nicorandil on HL-1 cardiomyocytes. PLoS One 12, e0172803 (2017).

62. Rawat, P. S., Jaiswal, A., Khurana, A., Bhatti, J. S. & Navik, U. Doxorubicin-induced cardiotoxicity: An update on the molecular mechanism and novel therapeutic strategies for effective management. Biomed. Pharmacother. 139, 111708 (2021).

63. Stringer, C., Wang, T., Michaelos, M. & Pachitariu, M. Cellpose: a generalist algorithm for cellular segmentation. Nat. Methods 18, 100–106 (2021).

64. Song, W., Wang, H. & Wu, Q. Atrial natriuretic peptide in cardiovascular biology and disease (NPPA). Gene 569, 1–6 (2015).

65. Man, J., Barnett, P. & Christoffels, V. M. Structure and function of the Nppa–Nppb cluster locus during heart development and disease. Cell. Mol. Life Sci. 75, 1435–1444 (2018).

66. Lewis, Y. E. et al. Localization of transcripts, translation, and degradation for spatiotemporal sarcomere maintenance. J. Mol. Cell. Cardiol. 116, 16–28 (2018).

67. Smalec, B. M. et al. Genome-wide quantification of RNA flow across subcellular compartments reveals determinants of the mammalian transcript life cycle. bioRxiv 2022.08.21.504696 (2022) doi:10.1101/2022.08.21.504696.

68. Sardão, V. A., Oliveira, P. J., Holy, J., Oliveira, C. R. & Wallace, K. B. Morphological alterations induced by doxorubicin on H9c2 myoblasts: nuclear, mitochondrial, and cytoskeletal targets. Cell Biol. Toxicol. 25, 227–243 (2009).

69. Meissner, M. et al. Moderate calcium channel dysfunction in adult mice with inducible cardiomyocyte-specific excision of the cacnb2 gene. J. Biol. Chem. 286, 15875–15882 (2011).

70. Badia-i-Mompel, P. et al. decoupleR: ensemble of computational methods to infer biological activities from omics data. Bioinformatics Advances 2, vbac016 (2022).

71. Cohen, J. A Coefficient of Agreement for Nominal Scales. Educ. Psychol. Meas. 20, 37–46 (1960).

72. Subramanian, A. et al. Gene set enrichment analysis: a knowledge-based approach for interpreting genome-wide expression profiles. Proc. Natl. Acad. Sci. U. S. A. 102, 15545–15550 (2005).

73. Barbie, D. A. et al. Systematic RNA interference reveals that oncogenic KRAS-driven cancers require TBK1. Nature 462, 108–112 (2009).

74. Xie, Z. et al. Gene Set Knowledge Discovery with Enrichr. Curr Protoc 1, e90 (2021).

75. Huang, H. et al. CTCF mediates dosage- and sequence-context-dependent transcriptional insulation by forming local chromatin domains. Nat. Genet. 53, 1064–1074 (2021).

76. Emanuel, G., seichhorn, Babcock, H., leonardosepulveda & timblosser. ZhuangLab/MERlin: MERlin v0.1.6. (2020). doi:10.5281/zenodo.3758540.

77. Lian, X. et al. Robust cardiomyocyte differentiation from human pluripotent stem cells via temporal modulation of canonical Wnt signaling. Proc. Natl. Acad. Sci. U. S. A. 109, E1848–57 (2012).

78. Frankish, A. et al. GENCODE reference annotation for the human and mouse genomes. Nucleic Acids Res. 47, D766–D773 (2019).

79. Yates, A. D., et al. Ensembl 2020. Nucleic Acids Res. 48, D682–D688 (2020).

80. Marçais, G. & Kingsford, C. A fast, lock-free approach for efficient parallel counting of occurrences of k-mers. Bioinformatics 27, 764–770 (2011).

81. Gans, J. D. & Wolinsky, M. Improved assay-dependent searching of nucleic acid sequence databases. Nucleic Acids Res. 36, e74 (2008).

82. Rodriguez, J. M., et al. APPRIS 2017: principal isoforms for multiple gene sets. Nucleic Acids Res. 46, D213–D217 (2018).

