## Supplementary Table 1 for "Bento: A toolkit for subcellular analysis of spatial transcriptomics data"

| **Categories** | **Features** |
| --- | --- |
| **Distance** | 1. Cell inner proximity: The average distance between all points within the cell to the cell boundary normalized by cell radius. Values closer to 0 denote farther from the cell boundary, values closer to 1 denote closer to the cell boundary. 2. Nucleus inner proximity: The average distance between all points within the nucleus to the nucleus boundary normalized by cell radius. Values closer to 0 denote farther from the nucleus boundary, values closer to 1 denote closer to the nucleus boundary. 3. Nucleus outer proximity: The average distance between all points within the cell and outside the nucleus to the nucleus boundary normalized by cell radius. Values closer to 0 denote farther from the nucleus boundary, values closer to 1 denote closer to the nucleus boundary. |
| **Symmetry** | 4. Cell inner asymmetry: The offset between the centroid of all points within the cell to the centroid of the cell boundary, normalized by cell radius. Values closer to 0 denote symmetry, values closer to 1 denote asymmetry. 5. Nucleus inner asymmetry: The offset between the centroid of all points within the nucleus to the centroid of the nucleus boundary, normalized by cell radius. Values closer to 0 denote symmetry, values closer to 1 denote asymmetry. 6. Nucleus outer asymmetry: The offset between the centroid of all points within the cell and outside the nucleus to the centroid of the nucleus boundary, normalized by cell radius. Values closer to 0 denote symmetry, values closer to 1 denote asymmetry. |
| **Dispersion** | 7. Point dispersion: The second moment of all points in a cell relative to the centroid of the total RNA signal. This value is normalized by the second moment of a uniform distribution within the cell boundary. 8. Nucleus dispersion: The second moment of all points in a cell relative to the centroid of the nucleus boundary. This value is normalized by the second moment of a uniform distribution within the cell boundary. |
| **Density** | 9. *L-function max: The max value of the L-function evaluated at r=[1,d], where d is half the cell’s maximum diameter. 10. *L-function max gradient: The max value of the gradient of the above L-function. 11. *L-function min gradient: The min value of the gradient of the above L-function. 12. *L monotony: The correlation of the L-function and r=[1,d]. 13. *L-function at d/2: The value of the L-function evaluated at ¼ of the maximum cell diameter.   *The L-function measures spatial clustering of a point pattern over an area of interest. |
